# *N*^6^-methyladenosine in poly(A) tails stabilize *VSG* transcripts

**DOI:** 10.1101/2020.01.30.925776

**Authors:** Idalio J. Viegas, Juan Pereira de Macedo, Mariana De Niz, João A. Rodrigues, Francisco Aresta-Branco, Samie R. Jaffrey, Luisa M. Figueiredo

## Abstract

RNA modifications are important regulators of gene expression. In *Trypanosoma brucei*, transcription is polycistronic and thus most regulation happens post-transcriptionally. *N*^6^-methyladenosine (m^6^A) has been detected in this parasite, but its function remains unknown. Here we show that ∼50% of the m^6^A is located in the poly(A) tail of the monoallelically expressed Variant Surface Glycoprotein (*VSG*) transcript. m^6^A residues are removed from the *VSG* poly(A) tail prior to deadenylation and mRNA degradation. Using genetic tools, we identified a 16-mer motif in the 3’UTR of *VSG* that acts as a cis-acting motif required for inclusion of m^6^A in the poly(A) tail. Removal of this motif from the *VSG* 3’ UTR results in poly(A) tails lacking m^6^A, rapid deadenylation and mRNA degradation. To our knowledge this is the first identification of an RNA modification in the poly(A) tail of any eukaryote, uncovering a novel post-transcriptional mechanism of gene regulation.

## Introduction

*Trypanosoma brucei* (*T. brucei*) is a protozoan unicellular parasite that causes lethal diseases in sub-Saharan Africa: sleeping sickness in humans and nagana in cattle^1^. In humans, the infection can last several months or years mostly because *T. brucei* escapes the immune system by periodically changing its variant surface glycoprotein (*VSG*)^2^. The *T. brucei* genome contains around 2000 antigenically distinct *VSG* genes, but only one *VSG* gene is actively transcribed at a given time. Transcriptionally silent *VSG* genes are switched on by homologous recombination into the BES or by transcriptional activation of a new BES^2^, resulting in parasites covered by ∼10 million identical copies of the *VSG* protein^3^.

VSG is essential for the survival of bloodstream form parasites. VSG is not only one of the most abundant proteins in *T. brucei*, but it is also the most abundant mRNA in bloodstream forms (4-11% of total mRNA)^4,5^. *VSG* mRNA abundance is a consequence of its unusual transcription by RNA polymerase I and its prolonged stability. The half-life of *VSG* mRNA has been estimated to range from 90-270 min, contrasting with the 12 min, on average, for other transcripts^6^. The basis for its unusually high stability is not known. It is thought to derive from the *VSG* 3’UTR, which contains two conserved motifs, a 9-mer and a 16-mer motif, found immediately upstream of the poly(A) tail. Mutational studies have shown that the 16-mer conserved motif is essential for *VSG* mRNA high abundance and stability^7^, even though its underlying mechanism is unknown.

VSG expression is highly regulated when the bloodstream form parasites undergo differentiation to the procyclic forms that proliferate in the insect vector^8^. The BES becomes transcriptionally silenced and VSG mRNA becomes unstable^9^, which results in rapid loss of *VSG* mRNA and replacement of the VSG coat protein by other surface proteins (reviewed in^10^). The mechanism by which *VSG* mRNA becomes unstable during differentiation remain unknown. The surface changes are accompanied by additional metabolic and morphological adaptations, which allow procyclic forms to survive in a different environment in the insect host^10^.

RNA modifications have been recently identified as important means of regulating gene expression. The most abundant internal modified nucleotide in eukaryotic mRNA is *N*^6^-methyladenosine (m^6^A)^11,12^, which is widespread across the human and mouse transcriptomes and is often found near stop codons and the 3’ untranslated regions of the mRNA encoded by multiple genes^13,14^. m^6^A is synthesized by a methyltransferase complex whose catalytic subunit, METTL3, methylates adenosine in a specific consensus motif. Demethylases responsible for removing m^6^A from mRNA have also been identified. m^6^A affects several aspects of RNA biology, for instance contributing to mRNA stability, mRNA translation, or affecting alternative polyadenylation site selection (reviewed in^15^).

Here we show that *N*^6^-methyladenosine is an RNA modification enriched in *T. brucei* mRNA. Importantly, this study revealed that m^6^A is present in mRNA poly(A) tails, and half of m^6^A is located in only one transcript (*VSG* mRNA). We identified a cis-acting element required for inclusion of m^6^A at the *VSG* poly(A) tail and, by genetically manipulating this motif, we showed that m^6^A blocks poly(A) deadenylation, hence promoting *VSG* mRNA stability. We provide the first evidence that poly(A) tails of mRNA can be methylated in eukaryotes, playing a regulatory role in the control of gene expression.

## Results

### m^6^A is present in *T. brucei* messenger RNA

To investigate if *T. brucei* RNA harbours modified nucleosides that could play a role in gene regulation, we used liquid chromatography-tandem mass spectrometry (LC-MS/MS) to detect possible modifications of RNA nucleosides. Thirty seven possible modified nucleosides were found in total RNA (Table S1), most of which had been previously detected in *T. brucei* RNA including Am, which is found in the mRNA cap structure, and m^3^C, m^5^C and Gm in tRNA and rRNA^16–18^. Mass-spectrometry analysis of mRNA revealed a clear peak that corresponded to m^6^A. Given the importance of this modification for RNA metabolism in other eukaryotes, we focused on this specific modification in *T. brucei*.

First, we tested if m^6^A was mainly present in mRNA, or if it was present in other type of RNA molecules (rRNA, tRNA and other non-coding RNAs). We also compared two different stages of the parasite life cycle, the mammalian bloodstream and the insect procyclic forms. An m^6^A nucleoside standard was used as a control. The chromatograms of the poly(A)-enriched fraction (mRNA) revealed a single peak corresponding to the 282→150 mass transition and an elution time of 10 min (Fig. 1A), confirming it represents m^6^A. The chromatograms of total RNA and poly(A)-depleted samples also contained a peak with an identical mass transition, but the elution time was much earlier (6.5 min), which likely reflects *N*^1^-methyladenosine (m^1^A), a modification with the same mass and earlier elution times, and is commonly found in rRNAs and tRNAs^19,20^. The m^6^A peak in poly(A)-depleted RNA was barely detectable, indicating that most (if not all) m^6^A is present in mRNA and absent from rRNA and tRNAs. Similar results were obtained in RNA fractions obtained from the procyclic form insect stage (Extended Data Fig. S1A).

**Figure 1.**
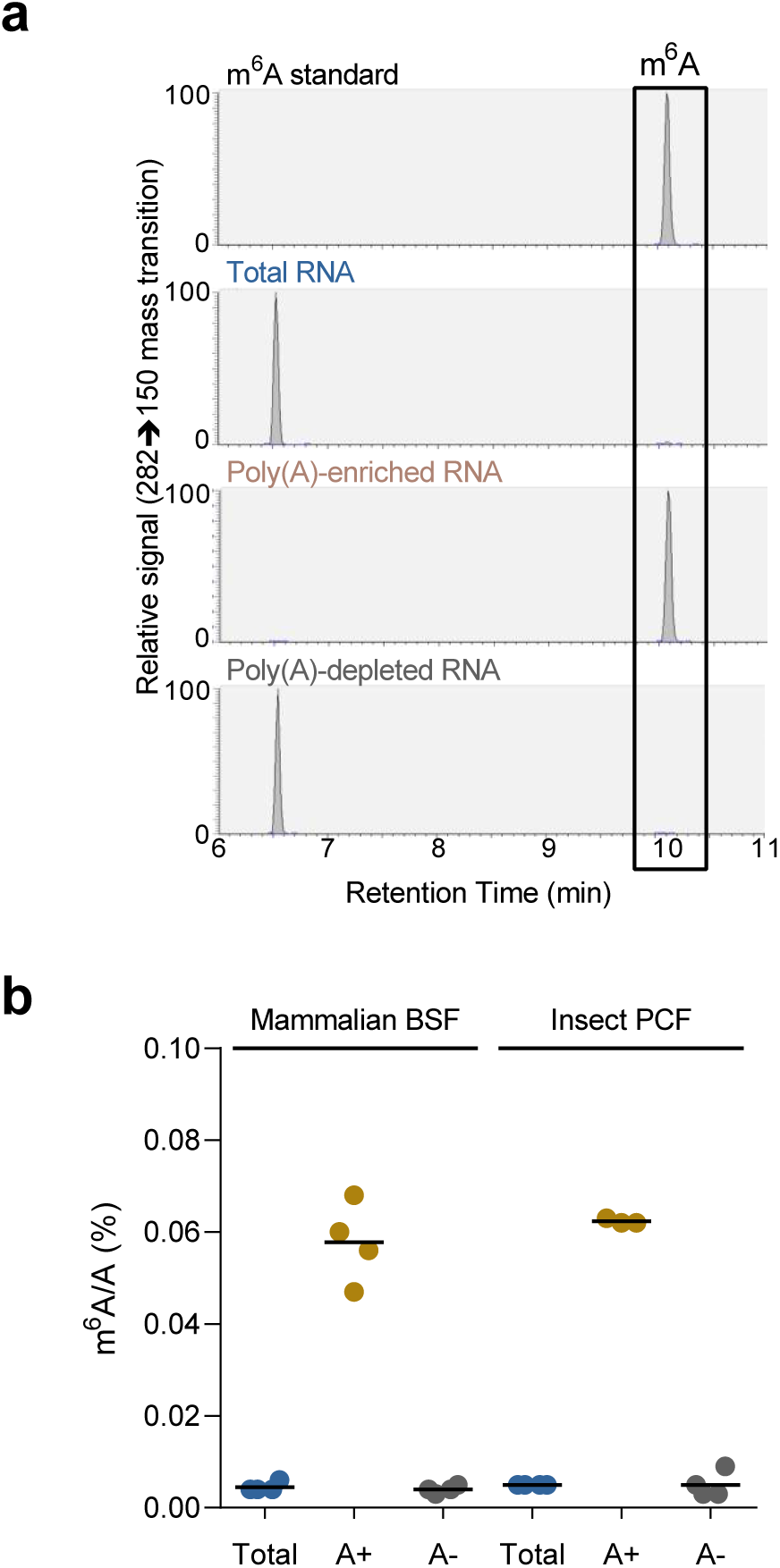
m^6^A is present in *T. brucei* mRNA in both insect and mammalian stages of the life cycle. **a**. Chromatograms obtained based on an LC-MS/MS analysis of a m^6^A standard and three RNA samples of *T. brucei* mammalian stages (BSF): total RNA, RNA enriched with oligo(dT)-beads (i.e., poly(A)-enriched RNA) and RNA that did not bind poly(T)-beads (i.e., poly(A)-depleted RNA). RNA was hydrolyzed and dephosphorylated and individual nucleosides were resolved by HPLC. m^6^A was readily detected in the poly(A)-enriched RNA. Another peak eluting at ∼6.6 min, corresponding to the tRNA and rRNA-enriched nucleoetide m^1^A, was detected in the poly(A)-depleted and total RNA samples. Identical analysis was performed in RNA from the insect life-cycle stage (PCF) – Fig. S1A. **b**. Levels of m^6^A detected by LC-MS/MS in the RNA samples indicated in panel (A) and quantified using the standard curve in Fig. S1B.(see also Extended Data Fig. S1)

The m^6^A standard allowed us to quantify the abundance of m^6^A in both stages of the life cycle (Extended Data Fig. S1B). As expected, the levels are very low in total and poly(A)-depleted fractions. In contrast, in both bloodstream and procyclic forms mRNA fractions, m^6^A represents 0.06% of total adenines in mRNA (Fig. 1B). In other words, 6 in 10,000 adenosines contain a methyl group at position 6. This proportion is lower than in mammalian cells (0.1-0.4%,^11,12^).

### m^6^A is enriched in the *VSG* poly(A) tail

During their life cycle, parasites need to adapt to living in different environments. The most recent studies have shown that around 30% of transcripts and around 33-40% of proteins are differentially expressed between these two life cycle stages^21,22^. It is possible that m^6^A is not equally distributed qualitatively and quantitatively in the transcripts of the two stages of the life cycle. To test this hypothesis, performed immunoblotting with an antibody that recognizes m^6^A. This antibody specifically recognizes m^6^A with minimal cross-reaction with unmodified adenosine or m^1^A (Extended Data Fig. S2). Mouse liver total RNA was used as a control. As expected^13^, the immunoblotting of liver total RNA revealed a smear in the entire lane, indicating that multiple RNA molecules of different sizes harbour m^6^A (Fig. 2A).

**Figure 2.**
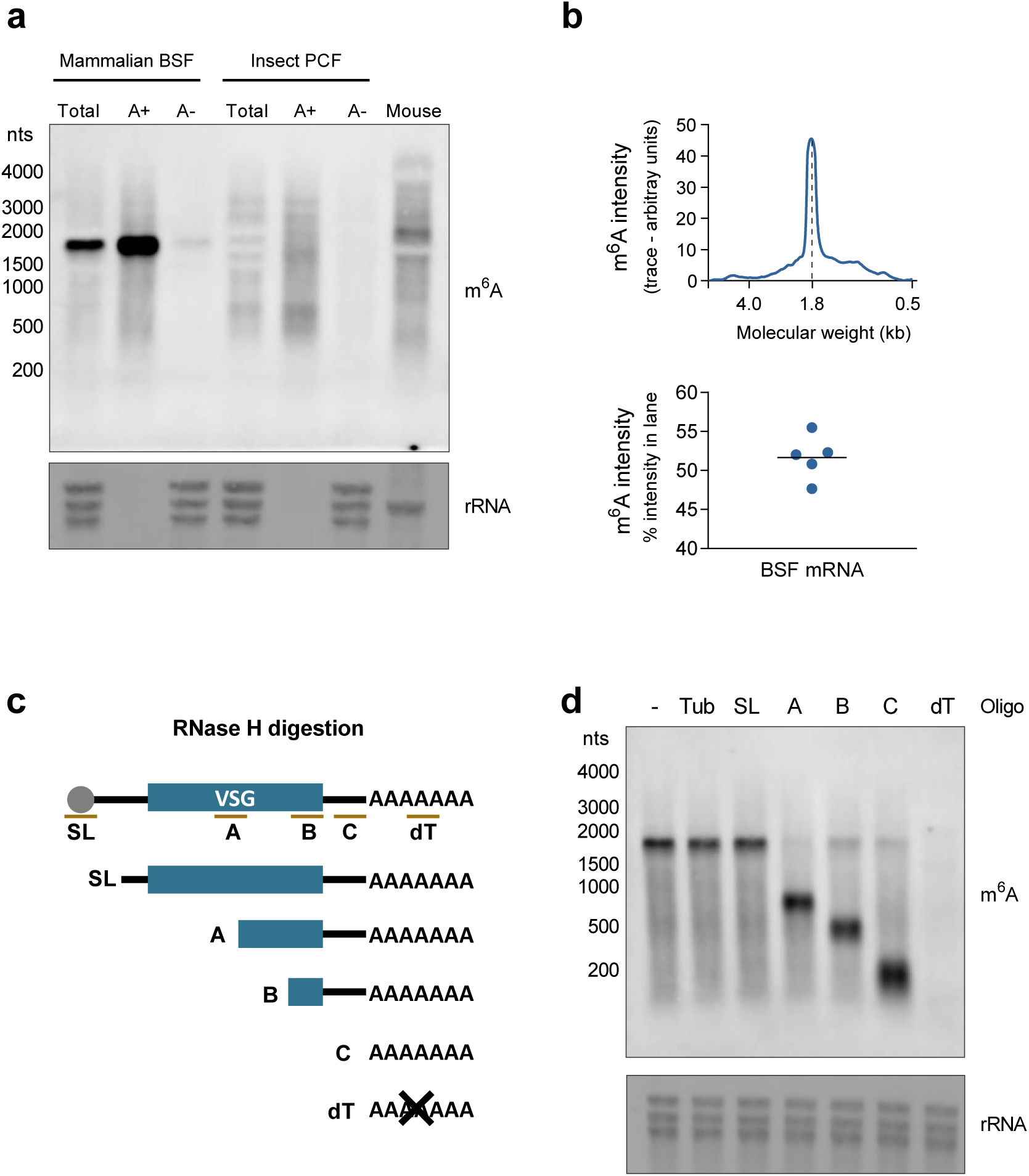
m^6^A is present in the poly(A) tail of *VSG* mRNA and other transcripts. **a**. Immunoblotting with anti-m^6^A antibody. RNA samples (from left to right): total RNA (Total), Poly(A)-enriched (A+) RNA and Poly(A)-depleted (A-) RNA from two life cycle stages (BSF and PCF). The last lane contains total mouse liver RNA (Mouse). 2 *µ*g of total RNA, 2 *µ*g of poly(A)-depleted RNA and 100 ng of poly(A)-enriched RNA was loaded per lane. rRNA was detected by staining RNA with methylene blue to confirm equal loading between total and poly(A)-depleted fractions. As expected the rRNA are undetectable in the poly(A)-enriched fraction. **b**. Intensity of the m^6^A signal in immunoblot, measured by Image J, in the whole lane containing the poly(A)-enriched RNA of bloodstream forms. The intensity of the ∼1.8 kb band was compared with the signal intensity of the entire lane, and averaged from 5 independent samples. **c**. Diagram displaying the location of the oligonucleotides used in RNase H digestion of VSG mRNA. The digestion products detected in the immunoblot (panel D) after incubation with each oligonucleotide (SL, A, B, C, dT) are also indicated. **d**. Immunoblotting with anti-m^6^A antibody of mammalian bloodstream forms total RNA pre-incubated with indicated oligonucleotides and digested with RNase H. 2 *µ*g of total RNA were loaded per lane. Staining of rRNA with Methylene Blue confirmed equal loading. SL: spliced leader, dT: poly deoxi-thymidines. Tub: α-Tubulin. (see also Extended Data Fig. S3)

Next, we examined m^6^A in total RNA, poly(A)-enriched and poly(A)-depleted RNAs derived from bloodstream and procyclic forms (Fig. 2A). Consistent with the results obtained by LC-MS/MS, the poly(A)-depleted fraction showed a weak m^6^A signal, revealing that m^6^A is absent or below detection levels in rRNAs and tRNAs. In contrast, an m^6^A-positive smear was detected in poly(A)-enriched fraction both in bloodstream and procyclic forms, confirming that m^6^A is present in multiple mRNA molecules in both stages of the life cycle. Strikingly, however, the sample of bloodstream forms revealed an intense band of around 1.8 kb. This band accounts for ∼50% of the m^6^A signal intensity in the lane (Fig. 2B), suggesting that m^6^A in bloodstream forms is highly enriched in one type or several similarly sized types of transcripts.

The most abundant transcript in bloodstream forms is the *VSG*, which accounts for ∼5% of the total mRNA ^5^. To test if the specific band is *VSG*, we repeated the immunoblot assay but site-selectively cleaved the *VSG* transcript with Ribonuclease H (RNase H) and monitored the mobility of the m^6^A band. Poly(A) RNA was incubated with DNA oligonucleotides that annealed at different sites along the length of the *VSG* transcript. RNase H digestion of the RNA:DNA hybrids result in fragments of predicted sizes (Fig. 2C-D). If the *VSG* transcript is the band with the intense m^6^A signal, the band detected by immunoblotting would “shift” to one or two of the fragments of smaller size.

We first performed RNase H digestion of RNA pre-incubated without any oligonucleotide or with a control oligonucleotide that annealed with α-tubulin (another abundant transcript in *T. brucei*) did not affect the mobility of the m^6^A band (Fig. 2D). Next, we used an oligonucleotide that hybridized to the spliced leader (SL) sequence, a 39nt sequence that contains the mRNA cap and that is added to every mRNA by a trans-splicing reaction^23^. RNase H digestion of RNA pre-incubated with the SL oligonucleotide showed no effect on m^6^A immunoblotting signal (Fig. 2D). This indicates that m^6^A is neither present in spliced leader sequence, nor in the mRNA cap structure. In contrast, when we used oligonucleotides *VSG*-A, *VSG*-B and *VSG*-C, which hybridized to three different unique sites in the *VSG* sequence (Fig. 2C), we observed that the major m^6^A band shifted, and in all three conditions, the 3’ end fragment contained the entire m^6^A signal. Importantly, *VSG*-C oligonucleotide is adjacent the beginning of the poly(A) tail. Thus, the 3’ fragment released upon RNase H digestion with *VSG*-C corresponds to the poly(A) tail of *VSG* mRNA. This fragment, which contains the entire m^6^A signal from the *VSG* transcript, is heterogeneous in length and shorter than 200 nt (Fig. 2D).

To further confirm that the 3’ fragment released after incubation with *VSG*-C and RNase H corresponds to the *VSG* poly(A) tail, we performed RNase H digestion in RNA pre-incubated with a poly(T) oligonucleotide. Consistent with the results using *VSG*-C, the major band detected by m^6^A-antibody completely disappears, further supporting the idea that in bloodstream forms most m^6^A is present in the poly(A) tail of *VSG* mRNA. Interestingly, digestion of RNA hybridized with poly(T), also abolished the smear detected by m^6^A-antibody, indicating that m^6^A present in non-*VSG* transcripts is also most likely located in their poly(A) tails. Notably, a similar approach to digest poly(A) tails does not affect m^6^A levels in mammalian mRNA of HeLa cells^13^. Thus, this is the first demonstration that a poly(A) tail can harbour a modified nucleotide.

Our results show that although *VSG* mRNA is only 4-11% of total mRNA, it accounts for ∼50% of cellular m^6^A, suggesting that *VSG* mRNA is preferentially enriched in m^6^A compared to other mRNAs. Based on the m^6^A frequency in the transcriptome and the enrichment in *VSG*, we estimate that there are nearly four m^6^A molecules per *VSG* mRNA. In contrast, among the non-*VSG* mRNAs, only one in five mRNAs are predicted to contain m^6^A.

### Removing m^6^A precedes deadenylation and degradation of *VSG* transcript

The half-life of *VSG* transcript is 90-270 min, while the median mRNA half-life in trypanosomes is 13 min^6^. Given that removal of the poly(A) tail often precedes RNA degradation, we hypothesized that the presence of m^6^A in the poly(A) tail could account for this exceptional *VSG* mRNA stability.

We tracked m^6^A levels in *VSG* mRNA as it undergoes degradation in three independent conditions. We first inhibited transcription in bloodstream form parasites with actinomycin D (ActD) and, for the next 6 hours, we quantified the amount of *VSG* mRNA that remains (by qRT-PCR). We also measured the levels of m^6^A in *VSG* (by immunoblotting) and the length of the *VSG* poly(A) tail using the Poly(A) Tail-Length Assay (PAT), which involves ligation of adaptors to the 3’ end of poly(A) tails and two consecutive PCRs using *VSG*-specific forward primers (Fig. 3A). The amplified fragments contain part of the ORF, the 3’UTR of *VSG* transcript and the downstream poly(A) tail, whose size is variable between different transcript molecules.

**Figure 3.**
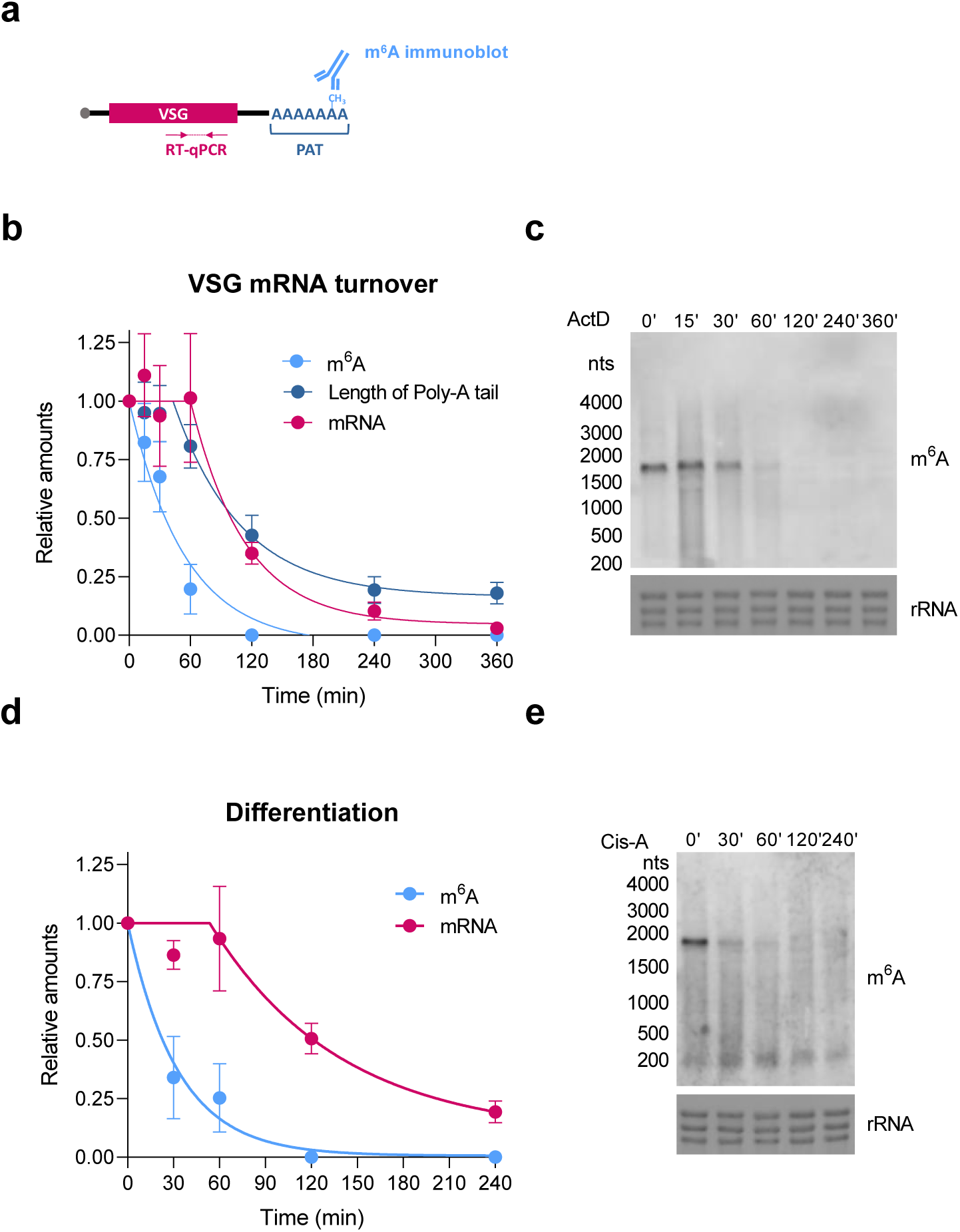
m^6^A is removed from *VSG* mRNA prior to its degradation. **a**. Diagram of *VSG2* mRNA transcript. Primers used for qRT-PCR are indicated in pink, primers used for PAT assay are indicated in dark blue and methylated adenosines of the poly(A) tail are indicated in light blue. **b**. Quantification of *VSG2* transcript levels, m^6^A signal and the length of poly(A) tail after transcription halt by actinomycinD (ActD). Total RNA of bloodstream parasites was extracted after various time-points of ActD treatment. Values were normalized to 0 hr. Transcript levels were measured by qRT-PCR (pink), m^6^A levels were measured by immunoblotting signal (light blue) and the length of the poly(A) tail was measured by PAT assay (dark blue). **c**. Immunoblotting with anti-m^6^A antibody of bloodstream form total RNA extracted from parasites treated with actinomycin D for increasing times. 2 *µ*g of total RNA were loaded per lane. Staining of rRNA with Methylene Blue confirmed equal loading. **d**. Quantification of *VSG2* transcript levels, m^6^A signal and the length of poly(A) tail during parasite differentiation from bloodstream to procyclic forms. Total RNA was extracted at different time-points after inducing differentiation with cis-aconitate. Values were normalized to 0 hr. Same color code as in Panel B. **e**. Immunoblotting with anti-m^6^A antibody of parasites differentiating to procyclic forms. 2 *µ*g of total RNA were loaded per lane. Staining of rRNA with methylene blue confirmed equal loading. (see also Extended Data Fig. S3)

*VSG* mRNA has previously been shown to exhibit biphasic decay: in the first hour after transcription blocking *VSG* mRNA levels remain high and, only in a second phase, do *VSG* mRNA levels decay exponentially^9,24^. Consistent with these earlier findings, we detected no major changes in mRNA abundance during the first hour after actinomycin D treatment (lag phase, or first phase); however, afterwards *VSG* exhibited exponential decay (second phase) (Fig. 3B). During the one hour lag phase, the length of the *VSG* poly(A) tail was unchanged, but then it rapidly shortened during the second phase. This indicates that there is a specific time-dependent step that triggers the rapid shortening of the *VSG* poly(A) tail and the subsequent degradation of the *VSG* transcript. Immunoblotting revealed that the m^6^A levels also decreased, but strikingly the loss of m^6^A preceded the shortening of the poly(A) tail and subsequent mRNA decay (Fig. 3C). In fact, m^6^A levels decrease exponentially during the first hour after actinomycin D, taking around 35 min for total mRNA m^6^A levels to drop 50%, while *VSG* mRNA only reached half of the steady-state levels around 2hr (Fig. 3B). These results indicate that m^6^A is removed from *VSG* mRNA prior to the deadenylation of the poly(A) tail, which is quickly and immediately followed by degradation of the transcript.

When bloodstream form parasites undergo cellular differentiation to procyclic forms, *VSG* is downregulated as a consequence of decreased transcription and decreased mRNA stabilty^9^. To test whether m^6^A is also rapidly removed from *VSG* mRNA prior to its developmentally programmed degradation, we induced differentiation *in vitro* by adding cis-aconitate to the medium and changing the temperature to 27°C. Parasites were collected and total RNA was extracted in different time points. Quantitative qRT-PCR showed that the levels of *VSG* mRNA stayed stable for around one hour, which was followed by an exponential decay (Fig. 3D). Importantly, immunoblotting analysis showed that during parasite differentiation, m^6^A intensity in the *VSG* mRNA dropped faster than the *VSG*-mRNA levels, reaching half of the steady-state levels in 23 min (Fig. 3D-E). Thus, during parasite differentiation, we observed again that the removal of m^6^A precedes the loss of *VSG* mRNA levels.

In addition to differentiation, another trigger for *VSG* mRNA degradation is translation inhibition, which can be achieved by treating parasites with drugs such as puromycin^25^. We treated bloodstream form parasites with puromycin and collected RNA samples for 4 hours. Consistent with previous studies, qRT-PCR revealed that *VSG* mRNA is actively degraded and it takes around 39 min to reach half of steady state levels (Extended Data Fig. S3). Immunoblotting revealed, once again, that m^6^A is removed earlier from the *VSG* transcript with a half-life of about 10 min (Extended Data Fig. S3).

Overall, these results show that in three independent conditions, m^6^A is removed from the *VSG* transcript earlier than the *VSG* transcript is deadenylated and degraded, suggesting that m^6^A may need to be removed from the *VSG* transcript before it can be degraded.

### Inclusion of m^6^A in *VSG* mRNA is dependent on *de novo* transcription

In most organisms, m^6^A is generated by methylation of adenosine residues within a specific consensus sequence by the METTL3 methyltransferase or its orthologs^15^. Given that in *T. brucei*, m^6^A is present in the poly(A) tail, a different mechanism is likely used. Indeed, trypanosomes lack a METTL3 ortholog^26^, indicating that a different pathway would be required to acquire m^6^A in the poly(A) tail. To understand how m^6^A accumulates in the VSG mRNA, we used parasite differentiation as a natural inducible system of *VSG* downregulation. This process is reversible in the first two hours^27^. Parasite differentiation was induced by adding cis-aconitate for 30 min (as described above, Fig. 3D), and then was washed away. As shown above, immunoblotting revealed that m^6^A was reduced after 30 min of cis-aconitate treatment (Fig. 4A-B, Fig. 3E). In contrast, when cells were allowed to recover for 1 hr in the absence of cis-aconitate or other drugs, we observed that the intensity of the m^6^A signal returned to normal levels (Fig. 4A-B). These results indicate that, if differentiation is halted and the levels of *VSG* mRNA are re-established (Extended Data Fig. S4), *VSG* m^6^A is quickly recovered.

**Figure 4.**
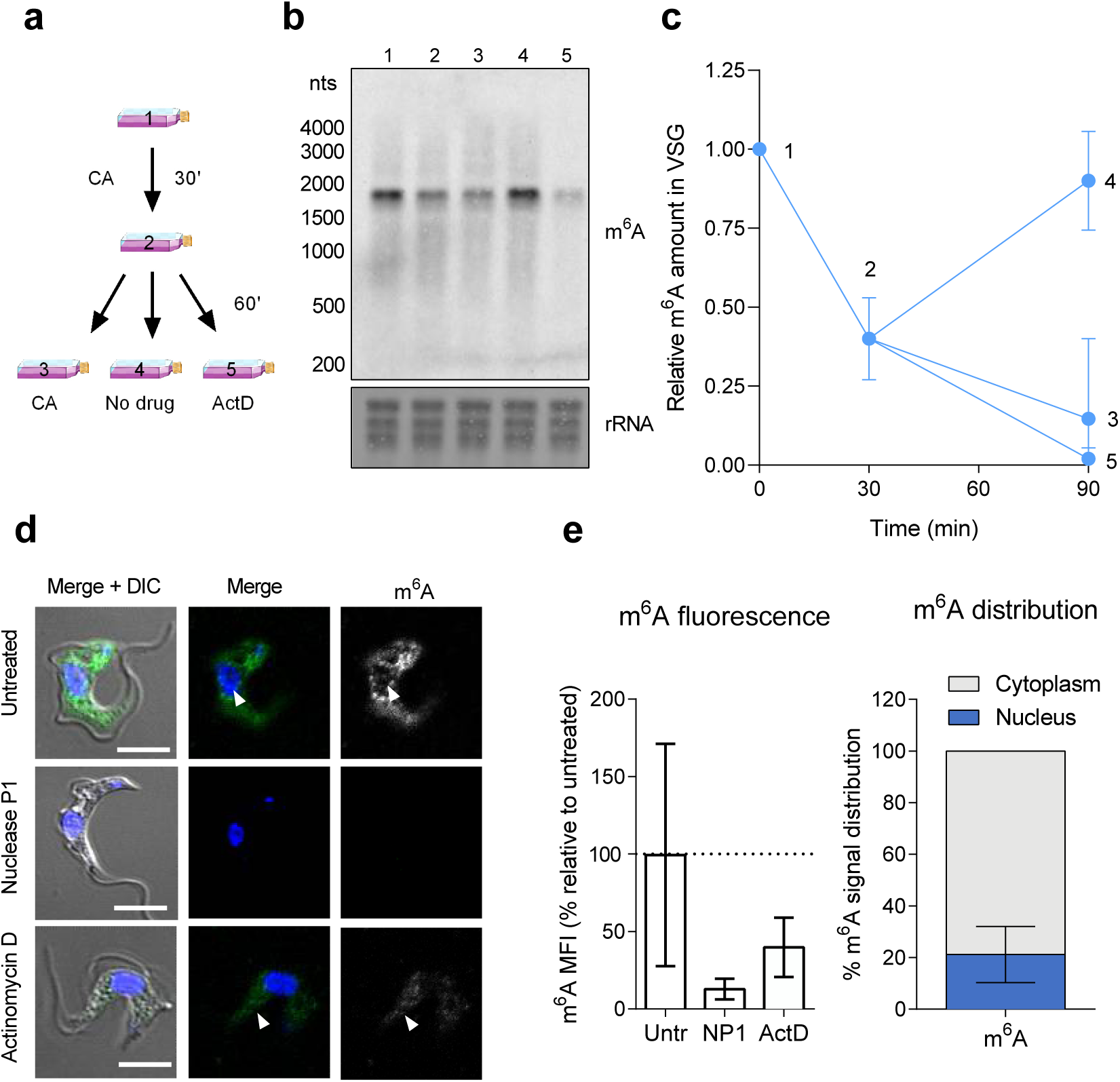
Inclusion of m^6^A in the *VSG* poly(A) tail depends of *de novo* transcription. **a**. Experimental setup. Parasites were treated with cis-aconitate (CA) for 30 min, time at which the compound was washed away. Parasites were placed back in culture in 3 different conditions for an extra hour: in the presence of cis-aconitate, without cis-aconitate, without cis-aconitate but with transcription inhibitor actinomycin D (ActD). Labels 1-5 indicate the conditions at which parasites were collected for Immunoblotting analysis (panel B). **b**. Immunoblot with anti-m^6^A antibody of total RNA from bloodstream parasites collected at each of the 5 labelled conditions (panel A). **c**. Quantification of Immunoblotting in panel A. All levels were normalized to time point 0 hr. **d**. Immunofluorescence analysis showing m^6^A localization in mammalian BSF. Parasites were treated with Nuclease P1 (NP1) or actinomycin D (ActD) to confirm signal-specificity. Nuclei were stained with Hoechst. Arrow in top row points to m^6^A signal in the nucleus. Arrow in bottom panel points to weak m^6^A signal; scale bars, 4*µ*m. **e**. Quantification of mean fluorescence intensity (MFI) levels of m^6^A in untreated BSF, nuclease P1 (NP1)-treated BSF, and actinomycin D (ActD)-treated BSF. Raw MFIs were obtained, the average of the untreated BSF equalled to 100%, and all other values normalized to 100%. Error bars indicate SEM. Results shown are the mean of *n* = 2 independent experiments, and *n* = 500 parasites per condition. **f**. Quantification of the proportion of m^6^A signal in nucleus and cytoplasm across 500 parasites (untreated BSF). The proportion was calculated by segmentation dividing the parasite body and nucleus, and quantifying fluorescence intensity in each segment. Error bars indicate SEM. DIC, differential interference contrast. (see also Extended Data Fig. S4)

To test if the recovery of m^6^A levels after cis-aconitate removal was due to *de novo* transcription, parasites were cultured in the presence of actinomycin D. We observed that the intensity of m^6^A in the*VSG* transcript was not recovered. Instead, the m^6^A levels decreased by ∼20% (Fig. 4A-B). Overall, these results indicate that *de novo* transcription is required to re-establish m^6^A levels in *VSG* mRNA.

If m^6^A is included in *VSG* mRNA soon after transcription, and if it remains in the poly(A) tail until it gets degraded, we should be able to detect m^6^A in the nucleus and in the cytoplasm of the parasites. Immunofluorescence analysis with an antibody against m^6^A showed a punctate pattern both in the nucleus (20%) and cytoplasm (80%) (Fig. 4D-E). To confirm that this m^6^A signal originated from RNA, we incubated the fixed cells with nuclease P1 prior to the antibody staining, which specifically cleaves single-stranded RNA without any sequence-specific requirement. This treatment caused a marked reduction in the intensity of the m^6^A signal, indicating that the immunoreactivity of the m^6^A antibody derives from RNA, and not non-specific interactions with cellular proteins (Fig. 4C-D). As an additional control, we treated with actinomycin D for 2 hr prior to immunofluorescence analysis. This treatment is expected to result in reduced cellular mRNA levels. Immunostaining with the m^6^A antibody showed a drop in the intensity of the m^6^A signal by around 40% (Fig. 4D-E), which is similar to the reduction observed by immunoblotting (Fig. 3B-C). Overall, these data support the idea that the m^6^A immunostaining results likely reflect m^6^A in mRNA and not a non-specific binding of the antibody to the cells.

### Inclusion of m^6^A in *VSG* poly(A) tail is dependent on neighboring cis-acting motif

m^6^A is added to the *VSG* mRNA poly(A) tail soon after transcription, probably still in the nucleus. m^6^A is not simply targeted to all poly(A) tails in the cell since, if this were the case, all mRNAs would be methylated and *VSG* methylation would only account for 4-11% of total cellular m^6^A. We therefore asked how the *VSG* poly(A) tail is selected for preferential enrichment of m^6^A. It has been previously shown that each *VSG* gene contains a conserved 16-mer motif (5’-TGATATATTTTAACAC-3’) in the 3’UTR adjacent to the poly(A) tail that is necessary for *VSG* mRNA stability^7^. It has been proposed that a currently unknown RNA-binding protein may bind this motif and stabilize the transcript by an unknown mechanism^7^. Here, we hypothesized that this 16-mer motif may act *in cis* to promote inclusion of m^6^A of the adjacent poly(A) tail.

To test the function of the 16-mer motif, we could not simply mutagenize it because this would lead to an irreversible loss of VSG protein, which is lethal for the parasites^7^. Therefore, we created two reporter cell-lines, in which a second *VSG* gene (*VSG117*) was introduced in the active BES, from where *VSG2* is normally transcribed (Fig. 5A). As a result, these reporter cell-lines are VSG double-expressors, i.e. they simultaneously express VSG2 and VSG117 at the cell surface. In the cell line designated “VSG double expressor 1”, or “DE1”, the *VSG117* gene contains the endogenous *VSG2* 3’UTR which harbors the 16-mer motif. In the second cell-line, “VSG double expressor 2”, or “DE2”, *VSG117* contains the same 3’UTR, except that the sequence of the 16-mer motif was scrambled (5’-GTTATACAAAACTTTT-3’) (Fig. 5A). As has been previously reported, the transcript levels of *VSG2* and *VSG117* are dependent on each other and are dependent on the presence of the 16-mer motif ^7^. Quantitative RT-PCR analysis showed that the two *VSG*s have roughly the same levels in DE1 cell-line. However, in DE2, *VSG117* transcript is about 7-fold less abundant than *VSG2*. (Fig. 5B, 5D).

**Figure 5.**
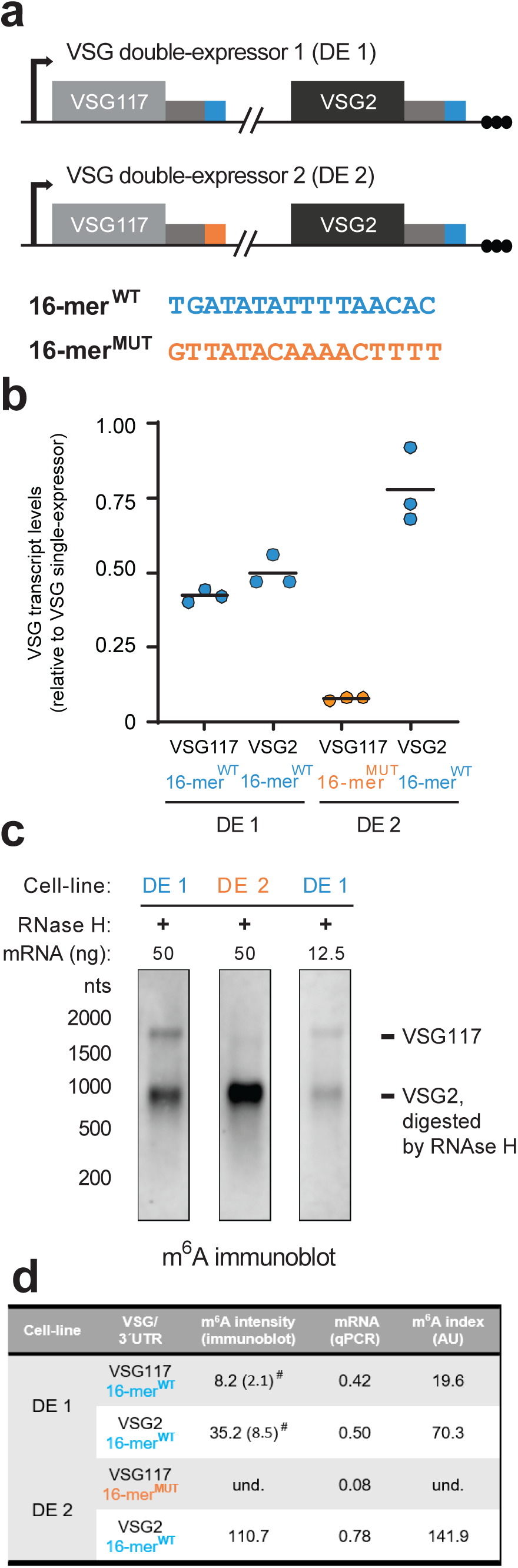
Conserved VSG 16-mer motif is required for inclusion of m6A in adjacent poly(A) tail. **a**. Diagram of 16-mer motif VSg double-expressor (DE) cell-lines. *VSG117* was inserted immediately downstream of the promoter of the active bloodstream expression site, which naturally contains *VSG*2 at the telomeric end. In VSG double expresor 16-mer^WT^ cell-line, *VSG117* contains its endogenous 3’UTR with the conserved 16-mer motif (sequence shown in blue). In VSG double expresor 16-mer^MUT^ cell-line, the 16-mer motif was scrambled (sequence shown in orange). **b**. Transcript levels of *VSG117* and *VSG2* transcripts measured by qRT-PCR in both reporter cell-lines. Levels were normalized to transcript levels in cell-lines expressing only *VSG2* or only *VSG117*. **c**. Immunoblot with anti-m^6^A antibody of mRNA from VSG double-expressor cell-lines. RNase H digestion of *VSG2* mRNA was used to resolve *VSG2* and *VSG117* transcripts. Different quantities of the same VSG double expresor 16-mer^WT^ cell-line was loaded in two separet lanes (50ng and 12.5ng) to show that the VSG117 band is detectable in both conditions. **d**. m^6^A index calculated as the ratio of m^6^A intensity and mRNA levels, measured in panels c. and b., respectively. und., undetectable. # intensities measured in lane 3 of Figure 5C (see also Extended Data Fig. S5)

To test whether the 16-mer motif is required for inclusion of m^6^A in *VSG* poly(A) tails, we performed m^6^A immunoblotting of cellular RNA obtained from the two double-expressor cell lines. Given that *VSG2* and *VSG117* transcripts have similar sizes (∼1.8 kb), we used RNase H to selectively cleave VSG2 before resolving the RNA on gel. *VSG2* cleavage was performed by incubating the total RNA sample with an oligonucleotide that hybridizes to the *VSG2* open reading frame, followed by incubation with RNase H (as described in Fig. 2). As expected, the *VSG2*-m^6^A-containing fragment is smaller and runs faster on gel (Fig. 5C). An “m^6^A index” was calculated by dividing the relative intensity of m^6^A in each *VSG* band (Fig. 5C) by the corresponding relative transcript levels measured by qRT-PCR (Fig. 5B). A low m^6^A index indicates a given transcript has fewer modified nucleotides (Fig. 5D).

Whenever the 3’UTR of *VSG* transcripts contain a 16-mer^WT^ (VSGs with a blue box in Fig. 5A), *VSG* m^6^A bands are detectable by immunoblot and the m^6^A index varies between 20-140 arbitrary units. In contrast, when the 16-mer motif is mutagenized (*VSG117* with orange box in Fig. 5A), the VSG m^6^A is undetectable (Fig. 5C), and the m^6^A index therefore is too low to calculate. These results indicate that the motif is required for inclusion of m^6^A in the VSG transcript.

In contrast, if the 16-mer motif played no role in m^6^A inclusion, the *VSG117* m^6^A index would be identical in both cell-lines (DE1 and DE2), i.e. around 20. Given that the qRT-PCR quantifications showed the relative intensity of *VSG117* 16-mer^MUT^ is ∼0.10 (Fig. 5B), the predicted intensity of the *VSG117* 16-mer^MUT^ m^6^A band would have been 20 × 0.10= 2.0 arbitrary units. To be sure that a band with this level of m^6^A would be detected on an immunoblot, we ran a more diluted DE1 RNA sample in lane 3 (Fig. 5C). The intensity of the *VSG117* 16-mer^WT^ band is 2.1 arbitrary units (Fig. 5C and 5D), and it is easily detected in the immunoblot. Given that we could not detect any band corresponding to a putative methylated *VSG117* 16-mer^MUT^ in DE2 (even after over exposure of the immunoblot, Extended Data Fig. S5), we conclude that the *VSG* conserved 16-mer motif is necessary for inclusion of m^6^A in the *VSG* poly(A) tail.

### m^6^A is required for *VSG* mRNA stability

The unusual localization of m^6^A in the poly(A) tail, and the lack of a METTL3 ortholog, suggests that the underlying biochemistry of m^6^A formation in trypanosomes is different from what has been described in other eukaryotes. Given that, at this stage the mechanism of m^6^A formation in the poly(A) tail is unknown and therefore cannot be directly blocked, we used the genetic mutants of the 16-mer conserved motif to enquire about the function of m^6^A in *VSG* mRNA.

To test the role of the 16-mer motif on poly(A) length on mRNA stability, we measured *VSG* mRNA stability in 16-mer^WT^ and 16-mer^MUT^ cell-lines. *VSG* mRNA half-life was measured by blocking transcription for 1 hr with actinomycin D (the duration of the lag phase during VSG mRNA decay) and the levels of *VSG* mRNA were followed by qRT-PCR. PAT assay clearly shows that, when the *VSG117* transcript contains the 16-mer motif (16-mer^WT^ cell-line), the length of the *VSG117* poly(A) tail is stable for 1 hr (Fig. 6A-B). In contrast, *VSG117* transcripts containing a scrambled 16-mer motif exhibited very rapid shortening of the poly(A) tail. In this case, there was no detectable lag phase—instead, the length of the poly(A) tail was reduced to 25% of its original length after just 15 min, and was undetectable after 1 hr (Fig. 6A-B). Consistent with the fast kinetics of poly(A) deadenylation, in the absence of an intact 16-mer motif, *VSG117* transcript levels decayed very rapidly with a half-life of ∼20 min (Fig. 6B).

**Figure 6.**
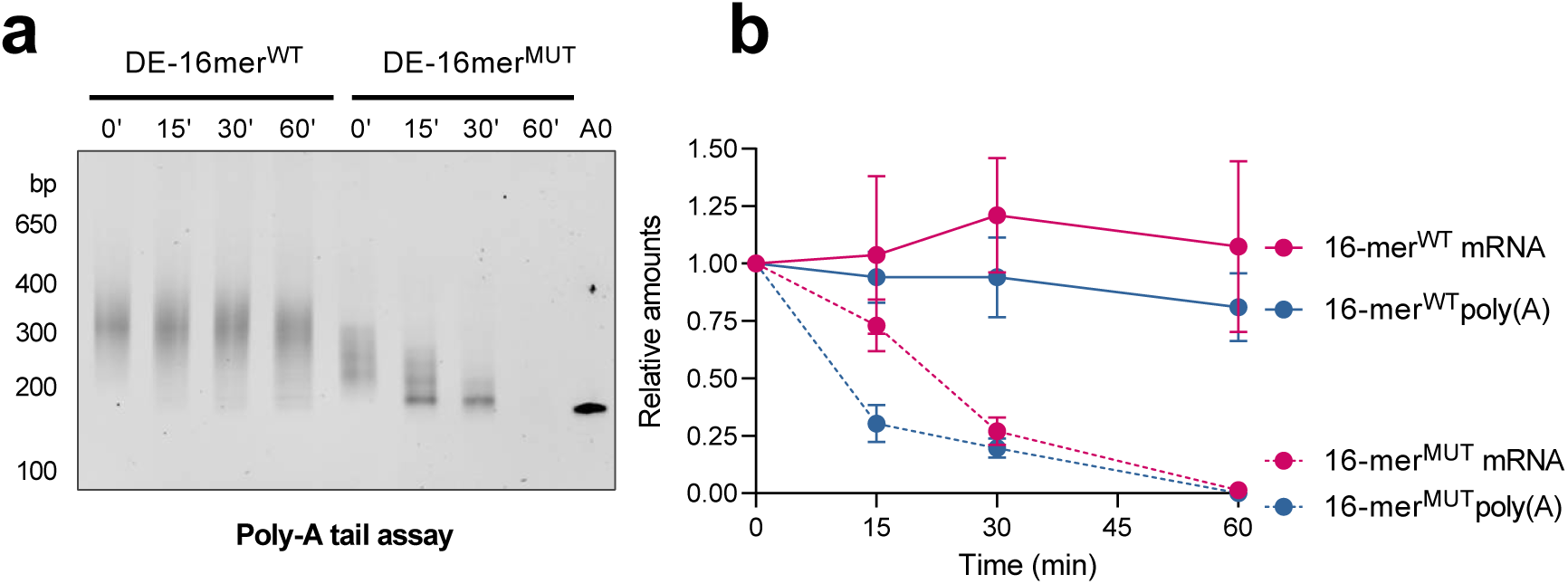
*VSG* 16-mer motif delays poly(A) tail deadenylation. **a**. The length of the *VSG* poly(A) tail was measured using Poly(A) tailing (PAT) assay. Specific primers confer specificity to the transcript being analysed. WT and Mut-16-mer cell-lines were compared. A0 corresponds to the minimum length amplified by PAT and it comprises the end of the *VSG* open reading frame and the whole 3’UTR. **b**. Quantification of *VSG117* transcript levels and the length of poly(A) tail after transcription halt by actinomycin D. Total RNA of bloodstream parasites was extracted after the indicate times. Values were normalized to 0 hr. Transcript levels were measured by qRT-PCR (pink) and the length of the poly(A) tail was measured by PAT assay (dark blue).

Overall, these experiments show that when the *VSG* conserved 16-mer motif is mutated and the m^6^A is lost, the *VSG* transcript is no longer stable and exhibits rapid poly(A) deadenylation and a marked reduction of mRNA stability.

## Discussion

In this work we report the identification and the function of the first RNA epitranscriptomic modification in the poly(A) tail of mRNA. We show that the stability and abundance of *VSG* transcripts stem from a novel mechanism in which the presence of m^6^A in the poly(A) tail inhibits RNA degradation, probably by blocking deadenylation. We also show that m^6^A is regulated by processes that induce destabilization and degradation of *VSG* mRNA such as cellular differentiation.

The classic function of a poly(A) tail is to keep mRNAs from being degraded and to promote translation. Poly(A)-binding proteins (PABPs) bind the poly(A) tail and stimulate mRNA translation by interaction with translation initiation factors^28^. Removal of the poly(A) tail by deadenylase complexes (e.g. Ccr4-Not complex) allow the unprotected mRNAs to enter in the 5’→3’ or 3’→5’degradation pathways^29,30^. In this report, we identified a novel mechanism by which a poly(A) tail contributes to mRNA stability. We found that the poly(A) tail of *VSG*, a transcript that represents 4-11% of mRNAs from *T. brucei* bloodstream forms, contains half of the m^6^A present in the cell. Our estimates suggest that each *VSG* poly(A) tail has around 4 modified adenosines. When we measured *VSG* mRNA decay rates by treating with transcription-blocking drugs, we found that m^6^A is removed from the *VSG* poly(A) tail before the tail is deadenylated and mRNA is degraded, raising the possibility that m^6^A normally prevents these processes. A similar observation was made during parasite differentiation. The removal of m^6^A may therefore be the step that initiates deadenylation and mRNA decay. To test this idea, we removed a 16-mer motif in the 3’UTR, which we find is necessary for the inclusion of m^6^A in the *VSG* poly(A) tail. When m^6^A levels in *VSG* mRNA become undetectable, by mutating this sequence element, the *VSG* mRNA exhibits rapid deadenylation and mRNA decay. Overall these data support the idea that m^6^A acts as a stabilizer of the *VSG* mRNA.

The presence of m^6^A in the poly(A) tail is so far unique to trypanosomes. In other eukaryotes, m^6^A has been mainly detected by m^6^A mapping approachs around the stop codon and 3’ UTR, where it plays a role in mRNA stability and translation^15^. A mapping study was recently published in *T. brucei* in which the authors conclude that m^6^A localizes in internal regions of transcripts^31^. m^6^A was not reported to be in the poly(A) tail in this previous study. However, m^6^A mapping relies on aligning m^6^A-containing RNA fragments to genomic sequence. Since the poly(A) sequence is not encoded in the genome, and m^6^A-containing fragment from the poly(A) tail would not be mappable and therefore not detected in this or any other previous m^6^A mapping study.

It remains unclear how m^6^A gets into the poly(A) tail. The presence of m^6^A in the poly(A) tail suggests that an unusual RNA-methyltransferase will directly or indirectly bind to the 16-mer motif and methylate adenosines that are either adjacent to the 16-mer motif or become more proximal via a loop-like conformation of the poly(A) tail. This would explain why orthologs of the canonical METTL3 enzyme does not exist in the trypanosome genome^26^. Unfortunately, previous efforts from other labs have been unsuccessful at identifying proteins that specifically bind to the 16-mer motif^7,32^. An alternative mechanism would be if m^6^A is incorporated into the poly(A) tail while polymerisation takes place. If this was true, there should be m^6^ATP inside the cell. Unbiased metabolomics^33^ did not support the existence of m^6^ATP, although studies specifically focused on detecting this nucleotide would be required to completely disprove this model.

Deadenylation is the first step in the main mRNA decay pathway in eukaryotes^34^. *T. brucei* is not an exception^23^. In this study we showed that m^6^A seems to protect the poly(A) tail from deadenylation. The molecular mechanism behind this stabilizing effect is unknown. It is possible that the deadenylases are inefficient on a methylated poly(A) tail. There is structural and biochemical evidence that poly(A) tails adopt a terciary structure that facilitates the recognition by some deadenylases (CAF1 and Pan2)^35^. When a poly(A) tail contains m^6^A, just like when it contains guanosines, the tertiary structure may be not properly assembled and deadenylase activity is inhibited. In this model, a putative demethylase may be required to remove the methyl group and only then poly(A) tail would be deadenylated. Alternatively, the stabilizing effect of m^6^A could result from recruitment of a specific RNA-binding protein, that prevents the poly(A) tail from being deadenylated.

*T. brucei* has around 2000 *VSG* genes, but only one is actively transcribed at a given time ^2^. By a process of antigenic variation, parasites evade the host immune response by changing an old VSG surface coat to a new VSG coat that is antigenically distinct. m^6^A may be removed from VSG mRNA, facilitating the switch from the old *VSG* mRNA, so that it can be more rapidly replaced by a new *VSG* coat protein. Since m^6^A stabilizes the *VSG* mRNA, it will also be important to determine whether inability to synthesize m^6^A in poly(A) tails results in more frequent switching to new *VSG* coat proteins. The role of m^6^A in differentiation remains to be established but it is possible that a delayed removal of *VSG* would interfere with the interaction of the parasite with the midgut wall.

The Rudenko lab has proposed that the maximal amount of *VSG* mRNA per cell is dependent on a post-transcriptional limiting factor dependent on the presence of the 16-mer motif ^7^. We propose that this limiting factor is the rate of inclusion of m^6^A in the poly(A) tails. When the 16-mer motif is present in both *VSG* genes, both get partially methylated and their abundance is reduced to about half of a single-*VSG* expressor; however, when the 16-mer motif is absent from one of the *VSG*s, the second *VSG* is more methylated and the transcripts become more abundant.

As far as we know, our work describes the first RNA modification in poly(A) tails. We show that m^6^A is present in the poly(A) tail of *T. brucei* mRNAs, it is enriched in the most abundant transcript (*VSG*), and that m^6^A acts as a protecting factor stabilizing *VSG* transcripts. It will be important for future studies to identify the enzymes and proteins involved in adding, reading, or removing this RNA modification. Given the importance of VSG regulation for chronic infection and parasite transmission, drug targeting such enzymes is expected to result in an important loss of parasite virulence. Understanding these regulatory epitranscriptomic processes may open up possibilities for developing therapeutic strategies to treat sleepiness sickness.

## Supplementary information

Figures S1 to S5

Tables S1 to S3

## Acknowledgments

The authors thank support from Howard Hughes Medical Institute International Early Career Scientist Program [55007419] and European Molecular Biology Organization Installation grant [2151]. This work was also partially supported by ONEIDA project (LISBOA-01-0145-FEDER-016417) co-funded by FEEI - “Fundos Europeus Estruturais e de Investimento” from “Programa Operacional Regional Lisboa 2020” and by national funds from FCT - “Fundação para a Ciência e a Tecnologia”. Researchers were funded by individual fellowships from Fundação para a Ciência e Tecnologia (PD/BD/105838/2014 to IJV, SFRH/BD/80718/2011 to FAB); a Novartis Foundation for Biomedical-Biological research to JPM; a Human Frontier Science Programme long term postdoctoral fellowship to MDN (LT000047/2019); the GlycoPar Marie Curie Initial Training Network (GA 608295) to JAR. LMF is an Investigator of the Fundação para a Ciência e Tecnologia. Publication of this work was also funded LISBOA-01-0145-FEDER-007391, project cofunded by FEDER, through POR Lisboa 2020 - Programa Operacional Regional de Lisboa, PORTUGAL 2020, and Fundação para a Ciência e a Tecnologia. We are grateful to Jane Thomas-Oates and Ed Bergstrom (York University, Centre of Excellence in Mass Spectrometry, Department of Chemistry) for the Mass-spectrometry analysis; Antonio Temudo, Ana Nascimento and Aida Lima for bio-imaging assistance; Ana Pena and members of Figueiredo and Jeffrey labs for helpful discussions.

## Author contributions

I.J.V., J.P.M., M.D.N and J.A.R performed experiments.

I.J.V., J.P.M., M.D.N, J.A.R, F.A.B, S.R.J and L.M.F planned experiments & analyzed data.

I.J.V., F.A.B., J.A.R., S.R.J and L.M.F conceived study.

I.J.V., S.R.J and L.M.F wrote manuscript, with contributions of all remaining authors.

## Methods

### Cell culture and cell-lines

*Trypanosoma brucei* bloodstream form (BSF) parasites (EATRO 1125 AnTat1.1 90:13, and Lister 427 antigenic type MiTat 1.2, clone 221a, Single Marker (SM) cell line^36^ were cultured in HMI11, supplemented with 10% Fetal Bovine Serum, at 37°C in 5% CO_2_^37^. Procyclic forms were cultured in DTM supplemented with 10% Fetal Bovine Serum at 27°C^38,39^. Parasites were routinely grown in the presence of the selectable drugs: neomycin 2.5 *µ*g·mL^-1^, hygromycin 5 *µ*g·mL^-1^ for EATRO 1125 AnTat1.1 90:13 and neomycin 2.5 *µ*g·mL^-1^ for SM. Transcription was inhibited by treating parasites with 5 *µ*g·mL^-1^ of actinomycin D (Sigma A4262). Differentiation of slender into stumpy forms was induced by adding 6 mM cis-aconitate (Sigma A3412) to the culture and dropping temperature to 27°C. Protein translation was inhibited with 50 *µ*g·mL^-1^ of puromycin (ant-pr-1, Invivogen).

Parasite cell lines were generated by transfection of *T. brucei* SM with plasmid p221-purVSG117UTR (Addgene plasmid 59732) or with p221-purVSG117UTRmut (Addgene plasmid 59732,^7^). The 16-mer mutagenized motif was introduced in a primer that was used to PCR amplify p221-purVSG117UTR plasmid. Amplification was performed with Phusion High-Fidelity DNA polymerase (ThermoScientific). After elimination of original plasmid template by digestion with *Dpn*I (NEB R0176), amplification products were transformed into *E. coli* JM109 (Promega L2005). Plasmids were isolated and purified from bacteria, digested with *Not*I-HF (NEB R3189) and *Xho*I (NEB R0146), ethanol precipitated and transfected. Transfections were made with the AMAXA nucleofector II (Lonza Bioscience), program X-001, using transfection buffer (90 mM sodium phosphate, 5 mM potassium chloride, 0.15 mM calcium chloride, 50 mM HEPES, pH 7.3). After overnight growth, transfected clones were selected by adding 1 *µ*g·mL^-1^ puromycin.

### LC-MS/MS

RNA samples were digested with nuclease P1 (sigma N8630, 1 U per 50 ng) for 2h at 37°C in buffer (2.5 mM ZcCl_2_, 40 mM NH_4_Ac, pH 5.3). After dephosporylation was done by addition of 10 U of Antarctic phosphatase (NEB M0289) in the buffer provided for 2h at 37°C. Digested samples were diluted in mili-Q water (10 ng/*µ*L) and filtered with Microcon – 30 kDa. Chromatographic separation was performed on a liquid chromatography systemUltimate 3000 RSLCnano (Thermo Fisher Scientific). Column – CORTECS® P3 (3 mm ×150 mm, 2.7 μm particle size, Waters Corporation). Mobile phase consisted of water containing 0.1% formic acid (A) and methanol containing 0.1% formic acid (B). The used elution gradient (A:B, v/v) was as follows: 100:0 for 2 min; 100:75 at 4 min; 75:30 at 9 min; 30:70 at 12 min; 0:100 at 15 min; 0:100 isocratic elution from 15.5 to 20 min. Samples were separated by liquid chromatography (column Waters CORTECS T3 2.7*µ* 3.0 × 150 mm, Gradient with mobile phase A of water 0,1% formic acid and mobile phase B of methanol 0,1% formic acid.) and analyzed in triple quadrupole Thermo Scientific TSQ Endura mass spectrometer with electrospray ionization in positive mode, and followed by multiple reaction monitoring. The ion mass transitions for m^6^A was 282,090>[150,059-150,061] and for adenosine was 268,061>[136,110-136,112]. Calibration curves were done with chemical standards and area under the curve (peak) integrated. To calculate m^6^A to adenosine molar ratio, m^6^A and adenosine amounts were calculated in the same sample injection. The ion mass transition of other modified nucleosides are shown in Supplementary Table 1.

### RNA isolation and handling

RNA was extracted from bloodstream and procyclic form parasites with TRIzol (Invitrogen). 10-100 million parasites were lysed in 1 mL of TRIzol. RNA was isolated according to the instructions of the manufacturer. RNA was treated with DNAse I (NEB M0303) (1 U per 2.5 *µ*g of RNA) for 20 min. Reaction was inactivated by adding 5 mM EDTA and heating to 75°C for 10 min. Purification of mRNA was performed with NEBNext mRNA isolation module (NEB E7490), following the manufacturer’s instructions. The RNA that did not bind to the poly(T)-beads was ethanol precipitated and saved. This RNA fraction was used as A-RNA.

cDNA was generated using a SuperScript II Reverse Transcriptase (Invitrogen 18064-014), using random hexamers, according to manufacturer’s protocol. Quantitative PCR (qPCR) was performed using SYBR Green PCR Master Mix (Applied Biosystems 4368702 Power SYBR). Primer efficiencies were determined using standard curves. Relative quantification was performed based on the CT (cycle threshold) value and the method of Pfaffl^40^. Amplifications were normalized to 18S ribosomal RNA transcript. Primers are indicated in Supplementary Table 2.

### m^6^A Immunoblotting

DNase I-treated RNA samples were suspended in formaldehyde loading buffer (30% formamide, 1,2 M formaldehyde, 1X MOPS buffer), denatured by heating to 70°C for 5 min and immediately transferred to ice. 2 □g of total RNA or 50 ng of mRNA were typically loaded per lane. Samples were resolved on a denaturing agarose gel (1.4% agarose, 2,2 M formaldehyde, 1X Mops buffer) for 1 h at 100 V at 4°C. RNA was transferred to a nylon membrane (GE Healthcare Amersham Hybond-N+) by downward transfer with 10X SSC buffer (1,5 M NaCl 150 mM sodium citrate, pH 7.0) for 4-5 h. RNA was UV-crosslinked to the membrane with a Stratalinker 2400 crosslinker (120 mJ cm^-2^). Membranes were stained with 0.02% methylene blue (Sigma M9140, diluted in 0,3 M sodium acetate pH 5.5) for 5 min and washed in RNase free water. After imaging, methylene blue was removed by incubation in de-staining solution (0,2X SSC 1% SDS) and washed 3 times in PBST (1X PBS pH 7.4 with 0,1% tween20). Nylon membranes were blocked by incubation in 5% skimmed milk in PBST for 1 h and then incubated overnight with 1*µ*g/mL rabbit anti-m^6^A antibody (Abcam ab151230, 1:1000 in 2.5% skimmed milk in PBST)at 4°C. Membranes were washed in PBST three times, 10 min each, and then incubated with HRP-conjugated donkey anti-rabbit IgG (GE Healthcare, NA934), diluted 1: 2500 in 2,5% skimmed milk in PBST for 1 h at room temperature. Membranes were washed in PBST, three times, 10 min each and signal developed with Western Lightning Plus-ECL, Enhanced Chemiluminescence Substrate kit (PerkinElmer ref. NEL103E001EA). The percentage of m^6^A in the main “band” co-migrating with the 2 kb molecular weight standard was measured by the intensity of the “band” divided by the signal intensity of the whole lane.

### Estimation of m^6^A per VSG mRNA

The number of m^6^A per VSG mRNA molecule was estimated based on the enrichment of m^6^A 1) in the VSG transcript (0,06%), 2) on the fraction of m^6^A present in *VSG* mRNA (0,50) and 3) on previously quantified parameters (detailed below). The number of mRNAs per *T. brucei* bloodstream forms cell is 20 000^41^. The number of VSG mRNAs per cell is 1000^41^ (assuming that correspond to 5% of the mRNA). The average mRNA length is 2000 nt^42^. The approximate length of the poly(A) tail is 100 nt^43^. The approximate frequency of adenosine in the transcriptome is 0,25.

> The number of adenosines per mRNA molecule correspond to:
>
> Number of adenosines (An) = (length of mRNA X frequency of adenosine) + poly(A) tail
>
> 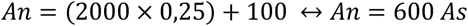
>
> The total number of adenosines in a bloodstream forms cell correspond to:
>
> An in bloodstream forms = mRNAs per cell X An per mRNA
>
> 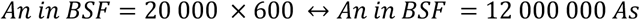
>
> The amount of m^6^A per cell in mRNA is:
>
> m^6^A in bloodstream forms = (adenosines per cell X Frequency of m^6^A in mRNA (%)) / 100
>
> 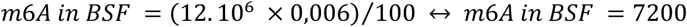
>
> The amount of m^6^A in *VSG* mRNA molecules correspond to:
>
> m^6^A in VSG mRNA = m^6^A in bloodstream forms X Fraction of m^6^A in VSG mRNA
>
> 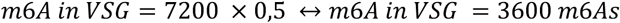
>
> The amount of m^6^A per VSG mRNA molecule correspond to:
>
> m^6^A per VSG = m^6^A in VSG/ VSG mRNA copies
>
> 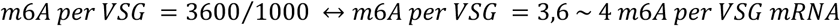
>
> Therefore, every *VSG* mRNA molecule harbors around 4 m^6^A per molecule.
>
> Following the same logic, the amount of m^6^A distributed in the non VSG mRNAs correspond to:
>
> m^6^A per non-VSG = m^6^A in non-VSG/ non-VSG mRNA copies
>
> 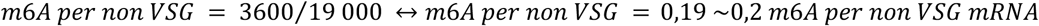
>
> Therefore, in the non-VSG transcriptome there are around 1 m^6^A per 5 mRNA molecules.

### RNA ligase mediated poly(A) tail assay (PAT assay)

5’ phosphorylated DNA oligo adaptor (500 pmol) was linked to the free hydroxyl group at the 3’-end of transcripts (1 □g of total RNA) by T4 RNA ligase 1 (NEB M0204) in the presence of 15% (w/v) PEG 8000 at 25°C for 4 h. Note that the adaptor oligo has a 3’-end dideoxyadenosine to avoid ligation between the adaptors. The ligated RNA (250 ng) was used to generate cDNA with the SuperScript II Reverse Transcriptase (Invitrogen 18064-014) according the manufacturer’s protocol using a primer complementary the adaptor. A VSG specific forward primer and an adaptor specific reverse primer were used for the subsequent PCR amplification using Phusion High-Fidelity DNA polymerase (ThermoScientific) with increased MgCl_2_ concentration (3.5 mM) and reduced extension temperature (60°C). The amplification products were diluted 1:500 and used as a template in a second amplification (nested PCR) with a second VSG specific forward primer and a second adaptor specific reverse primer. The A0 amplification product (Fig. 6) corresponds to the amplification product without a poly(A) tail. It results from a PCR amplification in which the reverse primer anneals at the end of the 3’ UTR immediately upstream of the poly(A) tail. Amplification products were resolved on a 6% TBE-PAGE (polyacrylamide gel electrophoresis) for 1 h at room temperature and stained with gel red for visualization. The length of the poly(A) tail estimated by co-migration of the amplification products with a DNA ladder minus the length of the A0 amplification product.

### Statistical analysis of decay curves

Data of the decay of mRNA, length of poly(A) tail and m^6^A signal were fitted in GraphPad Prism to either “Plateau followed by one phase decay” or “one phase decay”. The first were used when the variable decay started after an initial constant period. The “one phase decay” method was used when the measured variable decreased since the beginning of the experiment. The curves were adjusted to the data by the method of least squares, from which the decay constant K is estimated. The half-life was calculated by the ln(2)/K. This data is summarized in Supplementary Table 3.

### RNase H digestion

2 μg of total RNA was mixed with a complementary DNA oligo (0.2 pmol) in water and incubated 5 min at 70°C, after which temperature was reduced 1°C per min until 37°C. RNA:DNA hybrids were digested by RNase H (Ambion AM2293) for 30 minutes. The reaction was stopped by adding formaldehyde loading buffer and incubating at 70°C for 5 min. For RNase H digestions of *VSG2* transcript (Fig. 5), a thermostable RNase H (NEB M0523) was used to reduce unspecific digestion of the abundant *VSG117* transcript. Also mRNA, instead of total RNA, was used as the substrate to reduce the background and increase the sensitivity in the detection of *VSG117* transcript. The digestion mixture was prepared on ice with 50 ng of mRNA, (0.2 pmol) complementary DNA oligo, and Thermostable RNase H. The reaction mixture was transferred to the pre-warmed thermoblock at 50°C and incubated for 25 minutes.

### Immunofluorescence assays

Parasites were pelleted by centrifugation (800 g, 3 min, room temperature), resuspended in the remaining medium and transferred to an microcentrifuge tube. Pellet was resuspended in PBS and immediately fixed 4% v/v formaldehyde with gentle agitation by inversion. Fixed cells were centrifuged, resuspended in PBS and settled on poly-L-lysine coated coverslips for 1 h (for cells to settle by gravity). Parasites were permeabilized in 0,2% Nonidet P-40 in PBS for 5 min and washed in PBS three times for 5 min each. Cells were blocked in 1% BSA, 25 mg/mL glycine, in PBST for 1 h and then incubated with anti-m^6^A antibody (Abcam) 1: 250 (4 μg/ml^-1^ final concentration with 1% BSA in PBS) for 3 h at room temperature. After washing cells three times with PBS for 15 min each with agitation, cells were incubated with secondary anti-rabbit antibody (Alexa Fluor 488, A11034) for 30 min (protected from light). At the end, cells were washed three times with PBS for 15 min each and DNA stained for 1 min with 1 μg/ml Hoechst. Slides were mounted using Vectashield.

### Image acquisition and analysis

Confocal images on fixed *T. brucei* parasites were acquired using a Zeiss confocal Laser Point-Scanning (LSM) 880 Microscope equipped with the Zen 2.1 (black) software with a Plan Apochromat 63x NA 1.40 oil immersion DIC M27 Objective. The laser units used were a Diode 405-430 to excite the 405nm wavelength (corresponding to Hoechst), and an Argon laser to excite the 488nm wavelength (GaAsP detector 525/50 nm) (corresponding to AF488 used as a secondary antibody for m^6^A labelling). Images were acquired in a two-track mode. A pinhole diameter of 1AU for the 488 laser track was used. DIC images in confocal mode were obtained using the 405nm laser line. A digital zoom of 1.2x was used for general quantification, or a digital zoom of 4x for sub-localization analyses. For z-stack acquisition, on average 7-14 slices were obtained to cover all parasite areas per field, with a stack slice of 0.3*µ*m. Images were acquired in multiple fields of view to enable quantification of at least 100 parasites per condition per experiment repeat. Images were analysed using Fiji/ImageJ (imagej.nih.gov/ij/) for background correction, MFI determination and segmentation analysis. 3D rendering and 3D quantifications of m^6^A were performed using Imaris 9.1.0 (BitPlain). Statistical significance was determined using GraphPad Prism (GraphPad Prism Software version 6). Statistical tests used are mentioned in the corresponding Fig. legends. p<0.05 was considered significant.

