## Extended Data for "*N*^6^-methyladenosine in poly(A) tails stabilize *VSG* transcripts"

### SUPPLEMENTARY FIGURES and TABLES

A

B

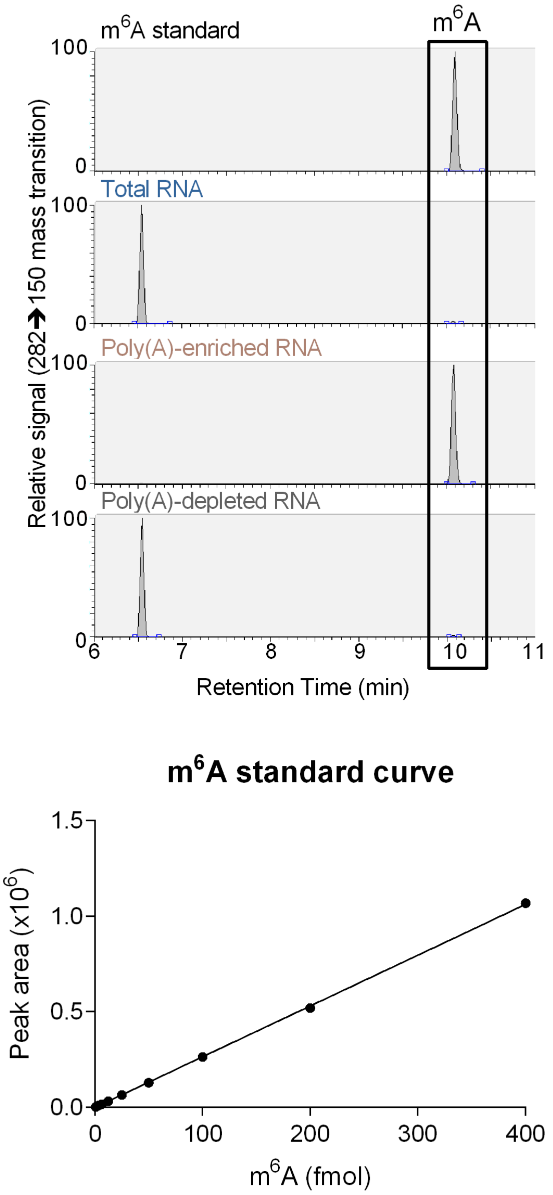


**Figure S1. Detection of m6A in insect stage (PCF) by mass-spectrometry.**
(**A**) Chromatograms obtained based on an LC-MS/MS analysis of a N6-methyladenosine standard and three RNA samples of *T. brucei* insect stages (PCF): total RNA, RNA enriched with poly(T)-beads (i.e., poly(A)-enriched RNA) and RNA that did not bind polyT-beads (i.e., poly(A)-depleted RNA). RNA was digested (nuclease P1 and antartic phosphatase) and individual nucleosides were resolved by HPLC and detected by mass spectrometry. Identical analysis was performed in RNA from the mammalian life-cycle stage (BSF) – Figure 2. (**B**) Standard curve of m$^6$A. Increasing quantities of commercially synthsized m$^6$A were loaded on the HPLC column and the area under the chromatogram peak was measured.

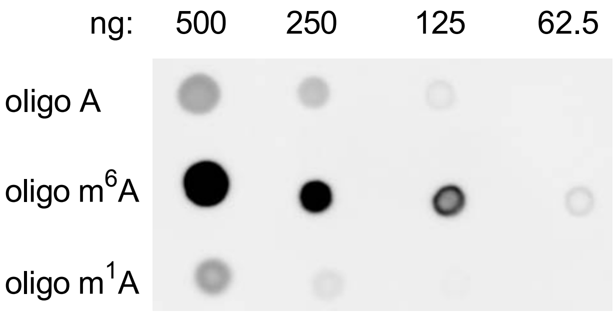


**Figure S2. Specificity of anti-m⁶A antibody.**
Oligonucleotides containing either m⁶A (positive control), unmodified adenosine and m¹A (negative controls) were manually spotted in the membrane, UV crosslinked and hybridized with anti-m⁶A anibody. The antibody specifically recognized the oligos with m⁶A, while exibiting low cross-reactivity to the oligos with only unmodified adenosine or containing m¹A.

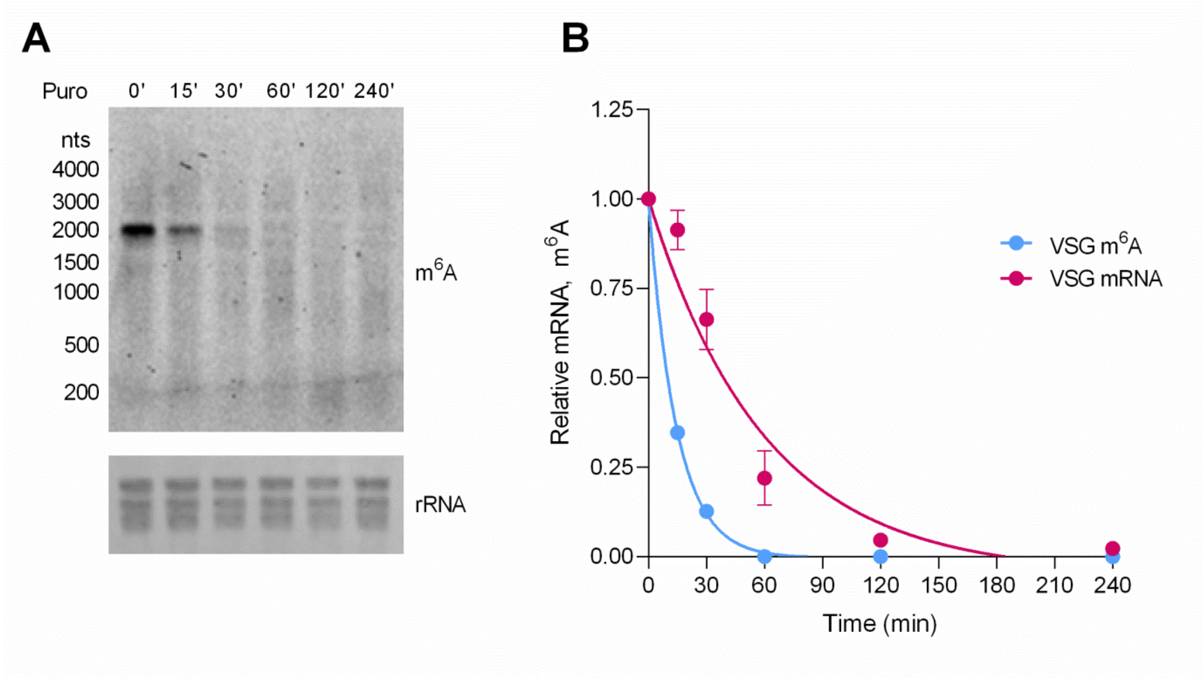


**Figure S3. Kinetics of VSG m⁶A and *VSG* mRNA upon translation blocking with puromycin.**
(**A**) Immunoblotting with anti-m⁶A antibody in parasites exposed to puromycin. Total RNA of bloodstream form parasites was extracted after various time-points of puromycin treatment. 2 μg of total RNA were resolved on gel. Loading was confirmed by staining rRNA with Methylene Blue. (**B**) Quantification of m⁶A (light blue) and *VSG* transcripts (pink) after translation halt by puromycin. Values were normalized to 0 hr. Transcript levels were measured by qRT-PCR and m⁶A levels were measured by immunoblotting.

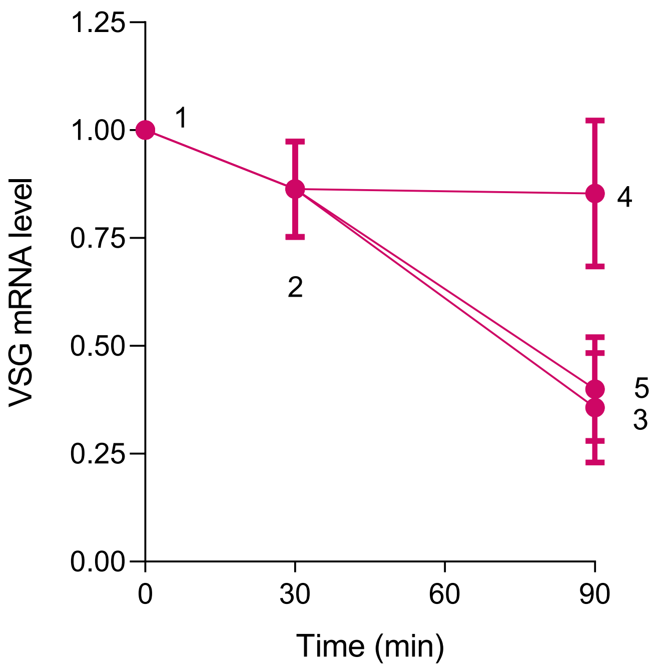


**Figure S4. qRT-PCR of samples analysed in Figure 4A-C.**
Total RNA was extracted from samples 1-5 indicated in Figure 4A. *VSG* mRNA levels were measured by RT-qPCR.

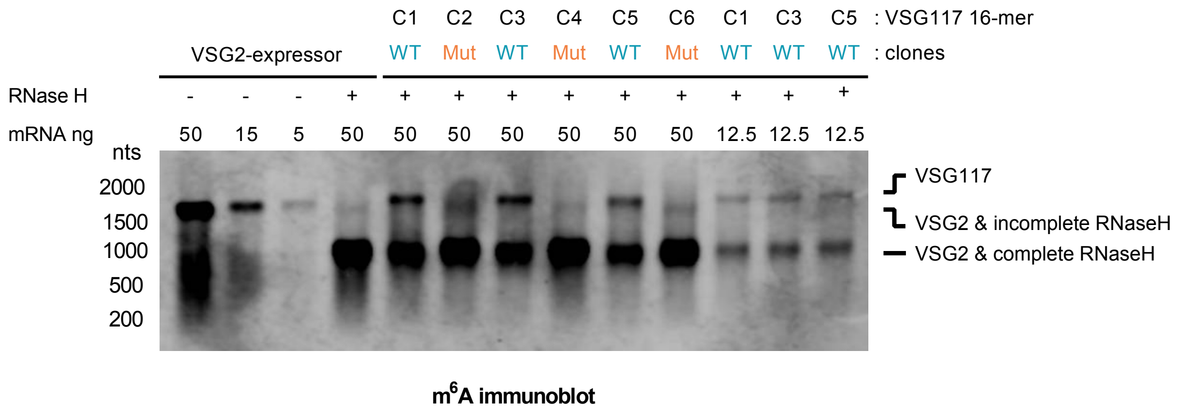


**Figure S5. Overexposure of immunoblot of Figure 5C.**
Overexposure of full immunobot presented in Figure 5C. Three independent 16-mer$^{WT}$ clones and three independent 16-mer$^{MUT}$ are shown (C1-C6). Note that with this exposure, most intense bands are saturated. The purpose of this hight exposure is to observe the region of blot corresponding to the *VSG117* transcript. No *VSG117* band is observed in the 16-mer$^{MUT}$ clones. It is also possible to observe a weak *VSG2* band in the *VSG2* single expressor lane and in the 16-mer$^{MUT}$ clones, which correspond to incomplete RNase H digestion of *VSG2* transcript.

**Table S1. 37 nucleoside modifications found in total RNA.**

| Base | Modification | Mass transition | Retention time (min) |
|---|---|---|---|
| A | $m^{1}A$ | 282→150 | 6.6 |
| | $m^{6}A$ | 282→150 | 10.0 |
| | $m^{6,6}A$ | 296→164 | 12.0 |
| | Am | 282→136 | 9.4 |
| | $m^{6,6}Am$ | 310→164 | 13.4 |
| | $i^{6}A$ | 336→204 | 16.9 |
| | $g^{6}A$ | 369→237 | 8.5 |
| | $ct^{6}A$ | 395→263 | 6.7 |
| | $t^{6}A$ | 413→281 | 12.9 |
| | $hn^{6}A/m^{6}t^{6}A$ | 427→295 | 13.2/14.1 |
| | $ms^{2}t^{6}A$ | 459→327 | 14.1 |
| C | $m^{3}C/m^{5}C/m^{4}C$ | 258→126 | 6.5/7.9 |
| | $m^{4,4}C$ | 272→140 | 7.9 |
| | Cm | 258→112 | 7.7 |
| G | $m^{7}G/m^{1}G/m^{2}G//hm^{6}A$ | 298→166 | 7.3/8.5/9.6/10 |
| | $m^{2,7}G$ /$m^{2,2}G$ | 312→180 | 8.5/10.8 |
| | $m^{2,2,7}G/ms^{2}m^{6}A$ | 328→196 | 5.6 |
| | Gm | 298→152 | 9.9 |
| | OHyW | 525→393 | 13.8 |
| | $o^{2}yW$ | 541→409 | 4.8 |
| U | Y | 245→125 | 4.5 |
| | $nm^{5}s^{2}U$ | 290→158 | 4.8 |
| | $ncm^{5}U$ | 302→170 | 7.9 |
| | $se^{2}U$ | 306→174 | 8.5 |
| | $mcm^{5}U$ | 317→185 | 7.9 |
| | ncm5s2U/nchm5U | 318→186 | 8.5 |
| | $cmnm^{5}U$ | 332→200 | 7.9/8.5 |
| | $acp^{3}U$ | 346→214 | 6.3 |
| | $cmnm^{5}s^{2}U$ | 348→216 | 3.8 |
| | $cmnm^{5}se^{2}U$ | 394→262 | 8.4 |
| | $nm^{5}ges^{2}U$ | 426→294 | 13.2 |

**Table S2. List of oligonucleotides.**

| Purpose | Sequence |
|---|---|
| AnTaT VSG qPCR | ACAACCACGGAAAGTGACG |
| AnTaT VSG qPCR | CACTTTTTGTCGCCATAAGC |
| VSG2 qPCR | AGCAGCCAAGAGGTAACAGC |
| VSG2 qPCR | CAACTGCAGCTTGCAAGGAA |
| VSG117 qPCR | AAGCGACAACAGATAAATGC |
| VSG117 qPCR | CTTTGCAAGCATTATTTTCC |
| 18S qPCR | ACGGAATGGCACCACAAGAC |
| 18S qPCR | GTCCGTTGACGGAATCAACC |
| 16-mer mutagenesis | TTTGTTATACAAAACTTTTCAAAACCAGCCGAGATTTTGTG |
| 16-mer mutagenesis | TTTGAAAAGTTTTGTATAACAAAAGTTTTCAAGTAGCAAGG |
| PAT adaptor | 5’-Ph-CCAGTGAGCAGAGTGACGAGGACTCGAGCTCAAGC-3ddA-3’ |
| PAT Rev_1 | GCTTGAGCTCGAGTCCTCG |
| PAT Rev_2 | CGTCACTCTGCTCACTGG |
| PAT rev A0 | GAGGACTCGAGCTCAAGCGCGTGTTAAAATATATCAG |
| PAT AnTaT 1 | AATCCCCGAATTGTAAATGG |
| PAT AnTaT 2 | TTTCTGCCGCATTTGTGG |
| PAT VSG117 1 | AAGCGACAACAGATAAATGC |
| PAT VSG117 2 | ATTCGCCCTCAGTGCTGC |
| AnTaT VSG RNase H A | TACTCGTCGTTGGCTGCTTG |
| AnTaT VSG RNase H B | TATTTTACTGCATAGGGCGT |
| AnTaT VSG RNase H C | GCGTGTTAAAATATATCAGA |
| Spliced leader RNase H | CAATATAGTACAGAAACTGT |
| β-tubulin RNase H | TACGGAGTCCATTGTACCTG |
| Oligo d(T) | TTTTTTTTTTTTTTTTTT |
| VSG2 RNase H | TCCGGCTGTTTCGTTTCT |
| Dot blot oligo A | ACTAGCTTAACTACGACCTCCTGAG |
| Dot blot oligo $m^6A$ | ACTAGCTTAACT-$m^6A$-CGACCTCCTGAG |
| Dot blot oligo $m^1A$ | ACTAGCTTAACT-$m^1A$-CGACCTCCTGAG |

**Table S3. Statistical parameters of time course experiments.**
Curves were fitted to the decay of *VSG* mRNA (pink), length of poly(A)-tail (dark blue) and m$^6$A levels (light blue). Curves in which the measured variable decayed from T=0hr were called "One phase". Those in which the measured variable decayed only after an initial constant period were called "Biphasic".

| Figure | Measure | Experimental approach | Curve fit | Lag phase (X0, min) | Decay constant (K, min$^{-1}$) | Half-life (min) | $R^2$ |
|---|---|---|---|---|---|---|---|
| 3B | mRNA | Transcription inhibition | Biphasic | 60.0 | 0.019 | 36.9 | 0.91 |
| 3B | poly(A) | Transcription inhibition | Biphasic | 42.9 | 0.015 | 45.4 | 0.95 |
| 3B | m$^6$A | Transcription inhibition | Exponential | NA | 0.019 | 37.4 | 0.93 |
| 3D | mRNA | Differentiation | Biphasic | 53.5 | 0.012 | 58.7 | 0.88 |
| 3D | poly(A) | Differentiation | Exponential | NA | 0.031 | 22.7 | 0.93 |
| 6B | mRNA | Transcription inhibition | Exponential | NA | 0.019 | 36.8 | 0.95 |
| 6B | poly(A) | Transcription inhibition | Exponential | NA | 0.074 | 9.3 | 0.98 |
| S3 | mRNA | Translation inhibition | Exponential | NA | 0.017 | 41.1 | 0.94 |
| S3 | m$^6$A | Translation inhibition | Exponential | NA | 0.070 | 9.9 | 0.99 |
